# CitriBEiTNet: A Hybrid CNN-Transformer Architecture Combining MobileNetV2 with BEiT’s Global Attention for Automated Citrus Leaf Disease Diagnosis

**DOI:** 10.64898/2025.12.09.693306

**Authors:** Hafza Eman, Syed M. Adnan, Wakeel Ahmad, Abid Ghaffar, Haider Ali Khan

## Abstract

Citrus farming plays an essential role in agriculture; however, diseases like canker, greening, black spot, and melanose significantly reduce yield and fruit quality. Efficient classification of citrus leaf diseases is important for crop health maintenance and optimal crop yield. Traditional methods for leaf disease detection are slow, labor-intensive, and often inaccurate, which highlights the need for automated solutions. This research presents a novel hybrid approach for identifying citrus diseases by combining a vision transformer with deep learning architectures. Using Bidirectional Encoder Representation from Image Transformers (BEIT) and MobileNetV2 as feature extractors, the proposed model captures distinctive features from images, which are then classified using Support Vector Machine (SVM). The dataset includes four different disease categories and a healthy class. Data augmentation techniques are applied to improve model robustness. The experimental findings demonstrate that CitriBEiTNet achieves a remarkable training accuracy of 99.82% and a testing accuracy of 99.57%, outperforming current leading techniques. This model provides an efficient, scalable, and economical approach for early disease identification, enabling farmers to take preventive measures and improve agricultural yields.

## Introduction

Agriculture is vital for the global economy, serving as the foundation for food production and sustaining the economic survival of millions of farmers globally. With the United Nations projecting an addition of 2 billion people to the global population within the next three decades, the demand for efficient and sustainable agricultural practices has never been greater [1]. As per the Food and Agriculture Organization (FAO), 691–783 million people experienced hunger in 2022, while close to 900 million suffered from severe food insecurity, highlighting the urgent need for sustainable agricultural solutions [2]. Since 2019, the number of undernourished people has risen by 180 million, highlighting an urgent need to enhance food production efficiency and crop resilience [2]. The increasing population in countries demands an increase in food as well [3]. The agriculture sector faces challenges due to crop losses caused by pests and diseases. They not only cause yield loss but also reduce the quality of crops. As a result, the economy and food supply chains are affected. Improving crop health and minimizing losses is important for sustaining agriculture and meeting the population’s needs [4].

Citrus fruits, including oranges, lemons, grapefruits, and mandarins, play an important role in global agriculture, contributing significantly to nutrition, economic stability, and trade. Citrus crops are important due to their high demand for fresh consumption and juice production worldwide [5]. Citrus is the most widely grown tree fruit globally, with an annual production of 104 million metric tons harvested across 7.1 million hectares of farmland [6]. Citrus production and exports have increased over the last thirty years [6]. In Pakistan alone, citrus farming accounts for nearly 30% of total fruit production, primarily concentrated in Punjab [7]. However, citrus cultivation faces several challenges, including weather conditions, pest infestations, disease outbreaks, limited availability of arable land, and technological limitations. One of the main challenges is that citrus plants suffer from many leaf diseases, such as canker, greening, black spot, and melanose, which cause harm to crops and the quality of products [8]. The excessive use of chemical pesticides to prevent disease damages the environment, harms wildlife, and affects human health. Early and precise identification of citrus diseases is essential to prevent disease outbreaks and maintain sustainable, high-quality fruit production [9].

Conventional disease diagnosis depends on visual expert assessments, a process that is inefficient, costly, and gives inconsistent results due to human subjectivity. The rapid development and refinement of computer vision and image processing in recent years have revolutionized plant disease detection, enabling faster, more accurate, and scalable identification methods [10]. A wide range of techniques, including deep learning, machine learning, and artificial neural networks, have been applied to automate the disease detection process [11]. Deep learning architectures have emerged as the superior approach for disease detection, due to their ability to automatically extract intricate visual features from vast and diverse datasets [12]. Modern AI systems using CNN and ViT architectures are achieving exceptional results in agricultural diagnostics, particularly for plant disease detection. While CNNs like DenseNet-201 and AlexNet have been applied to citrus disease classification with accuracies up to 99.6% [13], Transformer-based models such as BEIT remain underexplored in this domain.

To address these gaps, this research presents a new automated system that uses BEIT (Bidirectional Encoder Representation from Image Transformers) to accurately identify citrus diseases from leaf images. Our proposed framework utilizes BEIT’s unique bidirectional attention mechanism, which captures both local and global contextual features, which is a significant advantage over conventional CNNs that often miss long-range dependencies. The system incorporates an optimized feature extraction pipeline using BEIT’s hierarchical transformer architecture and an intelligent data augmentation strategy combining geometric transformations and sample generation. By enabling real-time, scalable disease detection, this technology empowers farmers to implement targeted interventions, reduce pesticide use, and optimize yields.

This paper is structured into five sections: Section 2 reviews current state-of-the-art methodologies, Section 3 describes the datasets used in this study and proposed methodology, Section 4 analyzes experimental findings through comparative results and ablation studies, and lastly, Section 5 concludes the work.

### Objectives

The objectives of this research are as follows:

- To develop an automated, hybrid model for classifying citrus leaf diseases.
- To achieve higher classification accuracy for multiple citrus diseases and healthy leaves in comparison with state-of-the-art methods.
- To evaluate the performance of each component in the model through ablation study.

## Literature Review

Plants are important for humans, serving as food and medicine for them. However, diseases that affect crops cause production loss and affect the economy. Accurate identification of plant diseases is essential for crop health. In the study [14], the comparison of Machine Learning (ML) and Deep Learning (DL) was presented. Support Vector Machine (SVM), Random Forest (RF), and Stochastic Gradient Descent (SGD) were used as Machine Learning models, such as Inception-v3, VGG-16, and VGG-19 were used as Deep Learning models. Deep learning methods perform better than machine learning methods in the case of disease detection as follows: Random Forest - 76.8% followed by Stochastic Gradient Descent - 86.5% followed by Support Vector Machine - 87% followed by VGG-19 - 87.4% followed by Inception-v3 - 89% followed by VGG-16 - 89.5%.

A two-stage deep CNN model for citrus disease classification is presented in [15]. The model consists of two stages. The first stage targets diseased areas on leaves, and the second stage is to classify the target area into the corresponding disease class. The proposed model achieved an accuracy of 94.37% and an average precision of 95.8%. A model was developed to detect and classify blackspot, canker, and greening diseases through images in [16]. The architecture used in this study has four blocks. Each block had a convolution, pooling, and batch normalization layer. The model achieved an accuracy of 96%.

In this study [17], the integrated approach is used, and a model is developed to distinguish between healthy and diseased leaves of a citrus plant. The diseased leaves are affected by common citrus diseases such as black spot, canker, scab, and greening. The model was tested on the citrus and plant village datasets. The model achieved an accuracy of 94.55%. A similar approach has been used by [18]. They used AlexNet and VGG19, two convolutional neural networks, and tested the proposed approach. The system’s performance reached 94% at its best. Another research [19] used two ways of conventional neural networks, named Alex Net and Res Net models. A self-dataset has been used for this research, which has 200 images of diseased and healthy citrus leaves. The trained models with data augmentation give the best results with 95.83% and 97.92% for Res Net and Alex Net, respectively.

The paper [20] used DenseNet 121 to identify diseases in citrus plants. The dataset used has a few citrus plant leaves divided into five categories. Four are diseased classes, and the fifth is healthy. The model, which is pre-trained on external data, achieved an accuracy of 92% and an F1 score of 95%. The combined pretraining model resulted in an accuracy of 88% and an F1 score of 88%. A new deep learning-based technique is presented by [21]. Two different pre-trained deep learning models have been used. Augmentation techniques have been used to increase the size of a dataset. The feature fusion technique is applied to combine features and then optimized using the Whale Optimization Algorithm (WOA). The model achieved an accuracy of 95.7%.

MobileNet and self-structured classifiers were used in [22] to detect and classify citrus leaf disease. The models were trained and evaluated on the citrus dataset, achieving a remarkable accuracy of 98% and 98%, respectively. The validation accuracies are 92% and 99%, respectively. [23] designed a model that is based on image processing and deep learning techniques. The dataset is divided into seven classes. The model achieved an accuracy of 90%. An Android application was also developed that used this model to diagnose disease. A hybrid approach has been used by [24] to identify citrus leaf diseases. The CNN model was used as a feature extractor, and the Random Forest was used as a classifier. The best model was VGG16-Random Forest, achieving an accuracy of 87%.

A patch-based classification network was proposed by [25]. The network consists of an embedding module, a cluster prototype module, and a neural network classifier. The model achieved an accuracy of 95.04%. The number of parameters was less than 2.3M, and the model is more efficient in terms of time in detecting disease. Bridge connections were used to improve the accuracy of models with some additional computational cost. The BridgeNet-19 was proposed for this study. The model achieved an accuracy of 95.47% [26]. The study introduced a unique method of identifying citrus leaf disease by combining CNN and Learning Vector Quantization (LVQ) techniques. Features are extracted through CNN, and LVQ is used to identify them as healthy or diseased. The model achieved an accuracy of 96.33% [27].

The study [28] proposed YOLO BP to detect green citrus in natural environments. CSPDarknet53 was used to extract features, and then Path Aggregation Network (PANet) and bi-directional feature pyramid network (Bi-PANet) were used to fuse the features. The results showed that the accuracy, recall, and mean average precision (mAP) of YOLO BP were 86, 91, and 91.55%. In another study [29] 500 images of vine leaves belonging to 5 classes were taken and then augmented to 2500 images were tested on the MobileNetv2 architecture. Features were extracted at the Logits layer, and classification was made using SVM kernels. 1000 features were extracted and then reduced to 250 using the Chi-Square method. The best SVM kernel was cubic, achieving an accuracy of 97.60

A pre-trained deep convolutional neural network is used in [30] as a feature extractor and a random forest as a classifier. The model achieved an accuracy of 91.66%. In [31], CNN models were used to identify diseases in citrus leaves. By using an ensemble strategy, the model was trained on varying numbers of classes. The study used leaf images and applied segmentation algorithms and feature extraction techniques. The model achieved an accuracy of 96%. A customized CNN-based model was developed by combining CNN with LSTM in [32]. The CNN-based method can differentiate between healthier fruits and leaves and diseased ones, such as fruit blight, fruit greening, fruit scab, and melanosis. The proposed model achieved 96% accuracy.

A CNN model was proposed in [33] that was trained on 596 images, and then augmented with 2800 images consisting of three categories. 70% of the data was used for training, 15% for validation, and 15% for testing. The training accuracy of the model was 95.95%, and the validation accuracy was 97.84%. The paper [34] presented a CNN-K that combines CNN with k-means clustering to segment images for identifying diseased areas and then classifying them. The hybrid model achieved an accuracy of 99.2% for all citrus fruit images. Although these studies have contributed a lot to plant disease detection, they still face some limitations. First, CNNs struggle with capturing long-range dependencies in images due to their localized receptive fields, which is problematic for analyzing disease symptoms that may be distributed across entire leaves. Second, they require substantial amounts of training data to achieve optimal performance, creating challenges for rare disease detection. Third, CNN architectures often incorporate significant inductive biases through their fixed filter designs, potentially limiting their ability to adapt to diverse disease patterns. Additionally, the hierarchical nature of CNNs may miss subtle but diagnostically important features that appear at multiple scales simultaneously.

BEIT addresses these limitations through its innovative architecture. BEIT has been successfully tested in several computer vision domains. The transformer-based architecture of BEIT offers several key advantages for plant disease detection. Firstly, its self-attention mechanism captures both local and global image features simultaneously. Secondly, it requires less training data than CNNs for comparable performance due to more efficient feature learning, and lastly, it can model relationships between spatially distant image regions that might collectively indicate disease presence. Furthermore, its pre-training on a massive dataset using a masked image modeling task allows it to develop a rich, general-purpose understanding of visual structures. This robust foundational knowledge can then be efficiently transferred to the specific task of identifying citrus diseases, even with limited field-specific imagery. These characteristics make BEIT particularly suitable for addressing the challenges in citrus disease detection, where early and accurate identification of complex symptom patterns is crucial for effective disease management.

## Experimental Methodology and Dataset

### Data Acquisition and Pre-processing

This study employs a citrus leaf disease dataset comprising 1,023 images sourced from Kaggle [10] and the PlantVillage dataset [35], featuring five distinct disease categories: black spot, canker, greening, melanose, and healthy. The study utilizes a Kaggle dataset containing 1,023 leaf images categorized into four groups: black spot (171 samples), canker (163), greening (204), and healthy specimens (485) shown in Figure 1. Before training, we applied pre-processing steps including size standardization, pixel normalization, and dataset splitting. Expert-validated class labels accompany each image to ensure reliable supervised learning. The Kaggle dataset is publicly available at: https://www.kaggle.com/datasets/sourabh2001/citrus-leaves-dataset/data.

**Fig 1.**
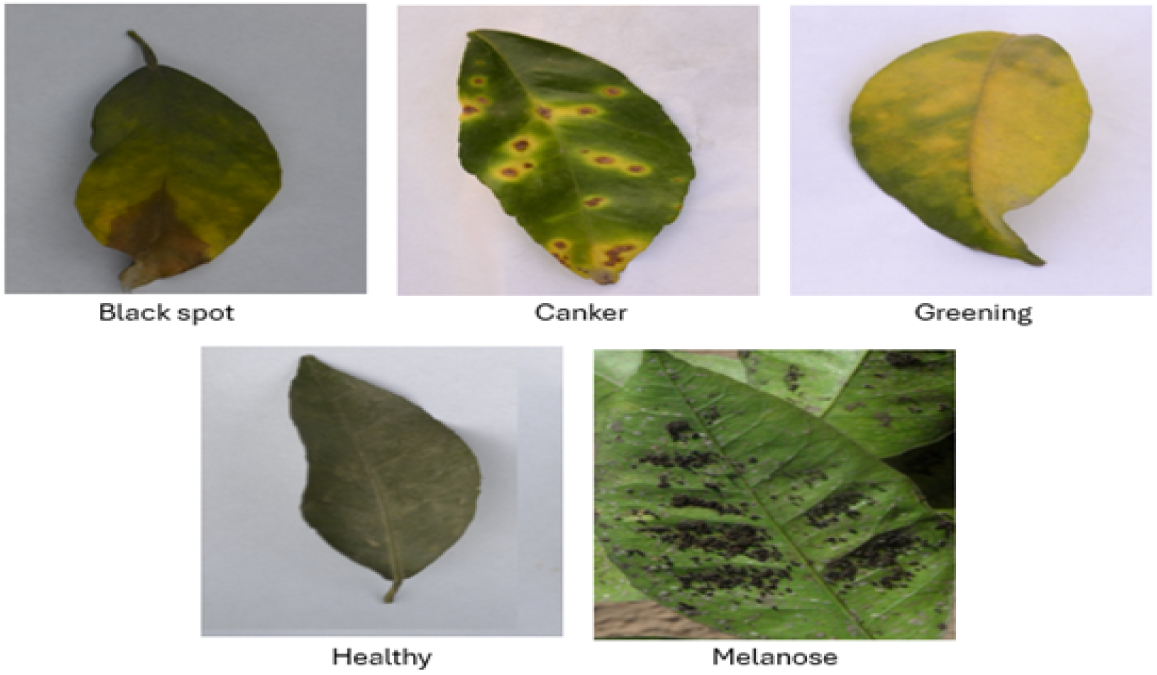
Sample images from the dataset showing leaf images from different disease categories.

To expand the training dataset and address class imbalance, artificial data generation techniques were systematically applied. This approach introduces controlled variations in the original images, enhancing model generalization while preventing overfitting through increased sample diversity. Rotational transformations included rotating the image left by 90 degrees, right by 270 degrees, and upside down by 180 degrees, simulating different orientations the object might appear in during real-world capture. Flipping operations were also performed, both horizontally and vertically, to ensure the model learns features invariant to reflection. Additionally, the brightness of the image was increased to simulate varying lighting conditions that can occur in natural environments. A geometric transformation in the form of a 15-degree shear was applied to mimic distortions caused by perspective changes or minor deformations. Collectively, these augmentations expanded the training data and supported the development of a more robust and reliable model. Figure 2 show augmentations applied to sample images from datasets.

**Fig 2.**
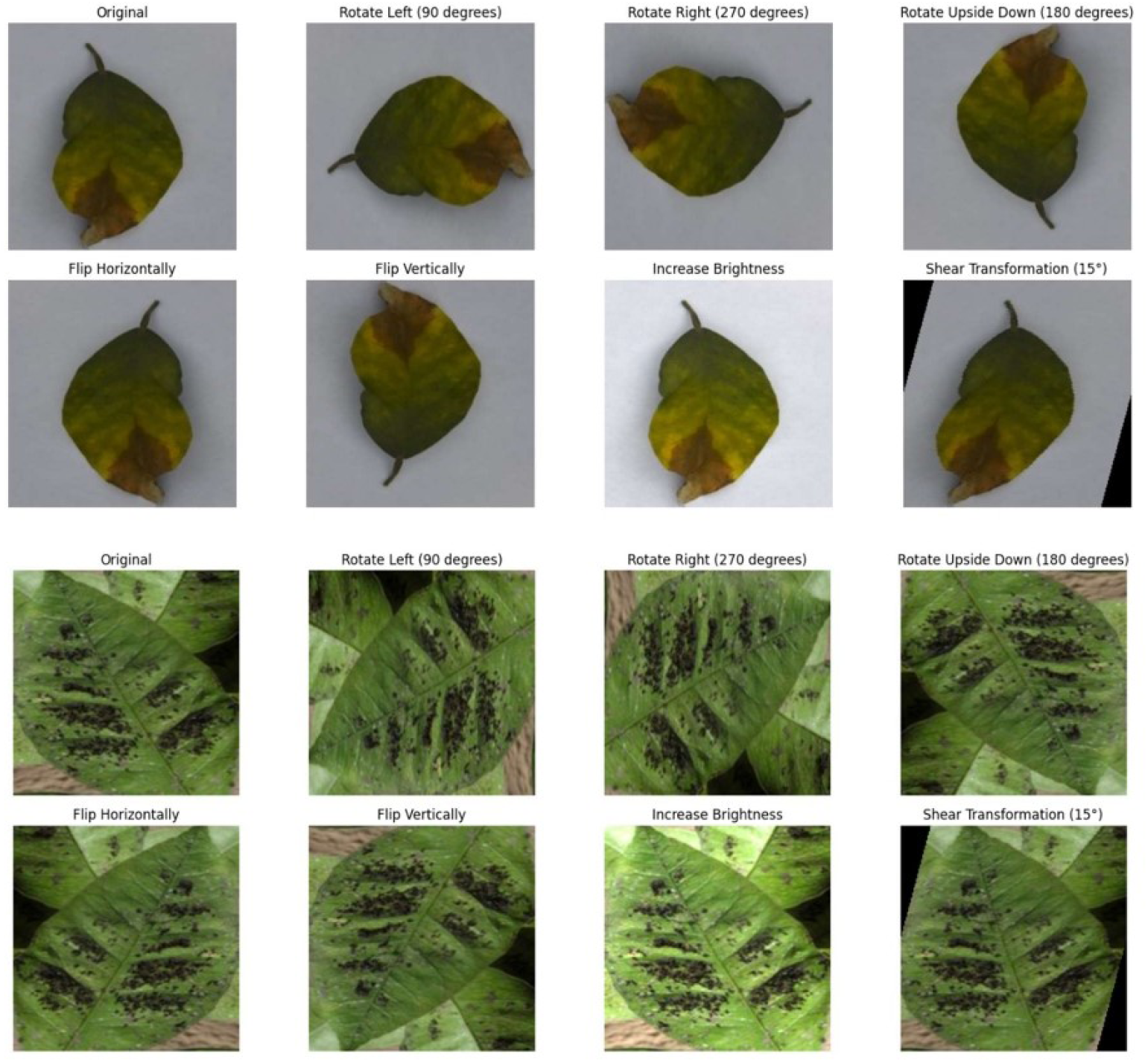
Visual demonstration of data augmentation techniques applied to citrus leaf images.

**Fig 3.**
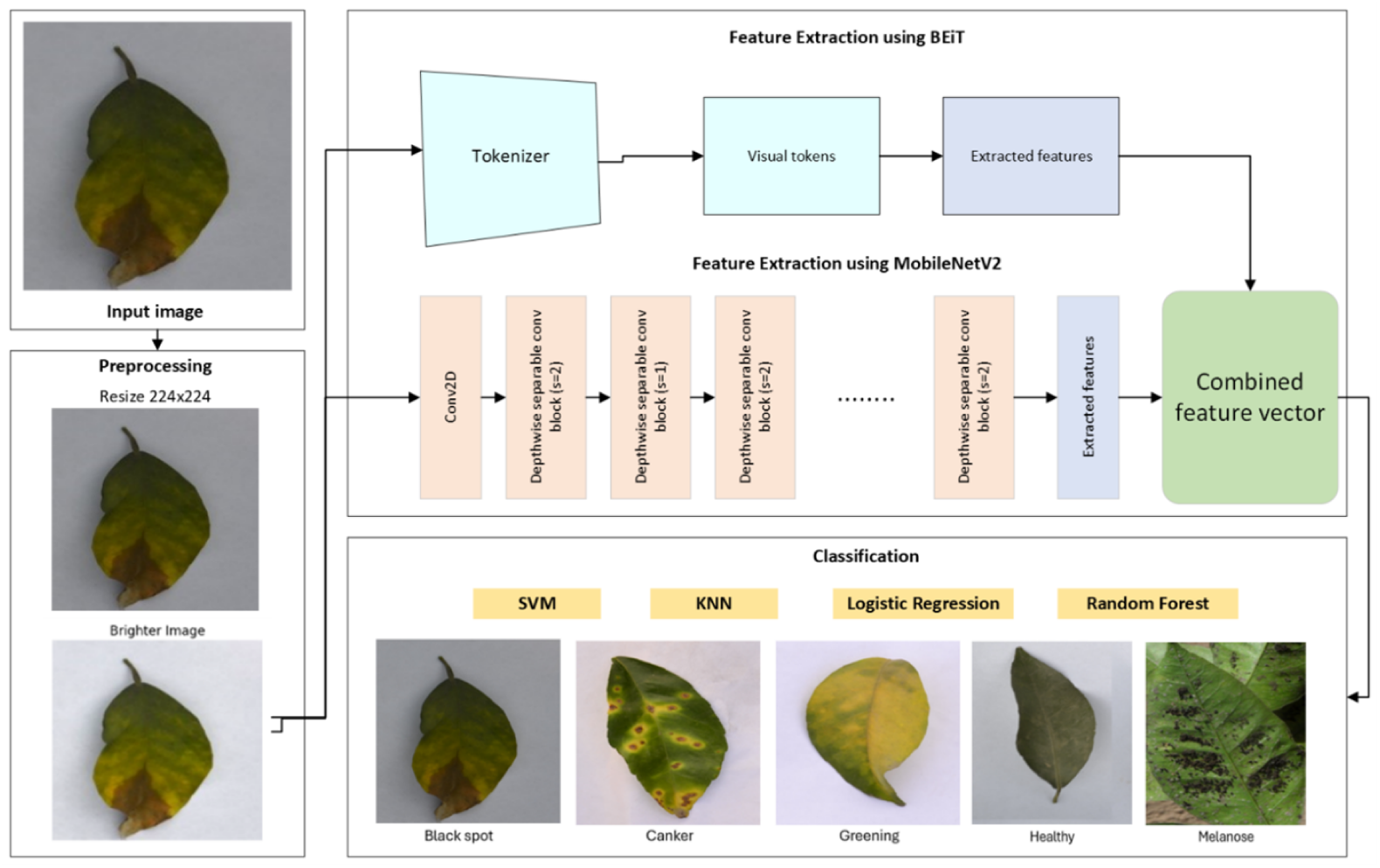
Architecture of the proposed hybrid pipeline for citrus disease classification, featuring BEiT and MobileNetV2 for feature extraction, with the resulting embeddings classified using Machine Learning classifiers.

### Methodology

The proposed methodology used a hybrid feature extraction and classification for leaf disease detection. Input images are preprocessed by resizing to 224×224 pixels and brightness level is also adjusted. Features are extracted using two models: BEiT (which tokenizes visual patches) and MobileNetV2 (which uses depthwise separable convolutions). The extracted features from both models are fused into a combined feature vector, which is then fed into traditional machine learning classifiers including SVM, KNN, Logistic Regression, and Random Forest to identify citrus leaf diseases.

### Feature Extraction Using BEiT

BEiT [36] is a vision transformer model introduced by Microsoft Research that adapts the principles of BERT (Bidirectional Encoder Representations from Transformers) to visual data. Unlike traditional CNNs, BEiT leverages a self-attention mechanism to model long-range dependencies between image patches, allowing it to learn rich contextual representations. It is pretrained in a self-supervised manner using a masked image modeling approach, where parts of the image are masked and the model learns to predict missing visual tokens, like masked language modeling in NLP.

The proposed system uses BEiT to extract useful features from images of citrus leaves for disease detection. The primary goal of using BEiT is to extract global, high-level features from citrus leaf images that capture the semantic structure of the visual data. These features can then be used for a classification task of identifying diseases in citrus leaves. Instead of relying on traditional methods that process images step by step with small filters, BEiT uses a transformer-based approach. This allows it to analyze relationships between different parts of the image, even if they are far apart. This is especially helpful for plant disease analysis because symptoms like spots, discoloration, or fungal growth can appear in irregular patterns across a leaf. By breaking the image into smaller patches and using self-attention mechanisms, BEiT can understand how diseased areas relate to healthy ones, leading to more accurate feature representations.

Citrus leaf images from datasets are resized and normalized using the BeitFeatureExtractor from HuggingFace’s transformer library. The extractor resizes images to 224 × 224 pixels (the input size BEiT expects) and standardizes pixel values using the ImageNet mean and standard deviation. BEiT then processes images by splitting them into fixed-size patches (e.g., 16×16), like tokens in NLP. For a 224×224 image, this results in 196 patches. These patches are linearly embedded and combined with positional embeddings before being passed through the transformer encoder [37]. The model used is Microsoft/Beit-base-patch16-224, which consists of 12 transformer layers, 12 attention heads, and a Hidden size of 768. The image is encoded through multiple self-attention layers [38] producing a sequence of embeddings, one for each patch, and an additional [CLS] token embedding that serves as a holistic representation of the entire image. The [CLS] token is a special token prepended to the input sequence. After the image passes through the transformer, the final state of the [CLS] token (a 768-dimensional vector) is extracted. This vector is considered the global feature representation of the image. Figure 4 shows a step-by-step feature extraction process using BEiT.

**Fig 4.**
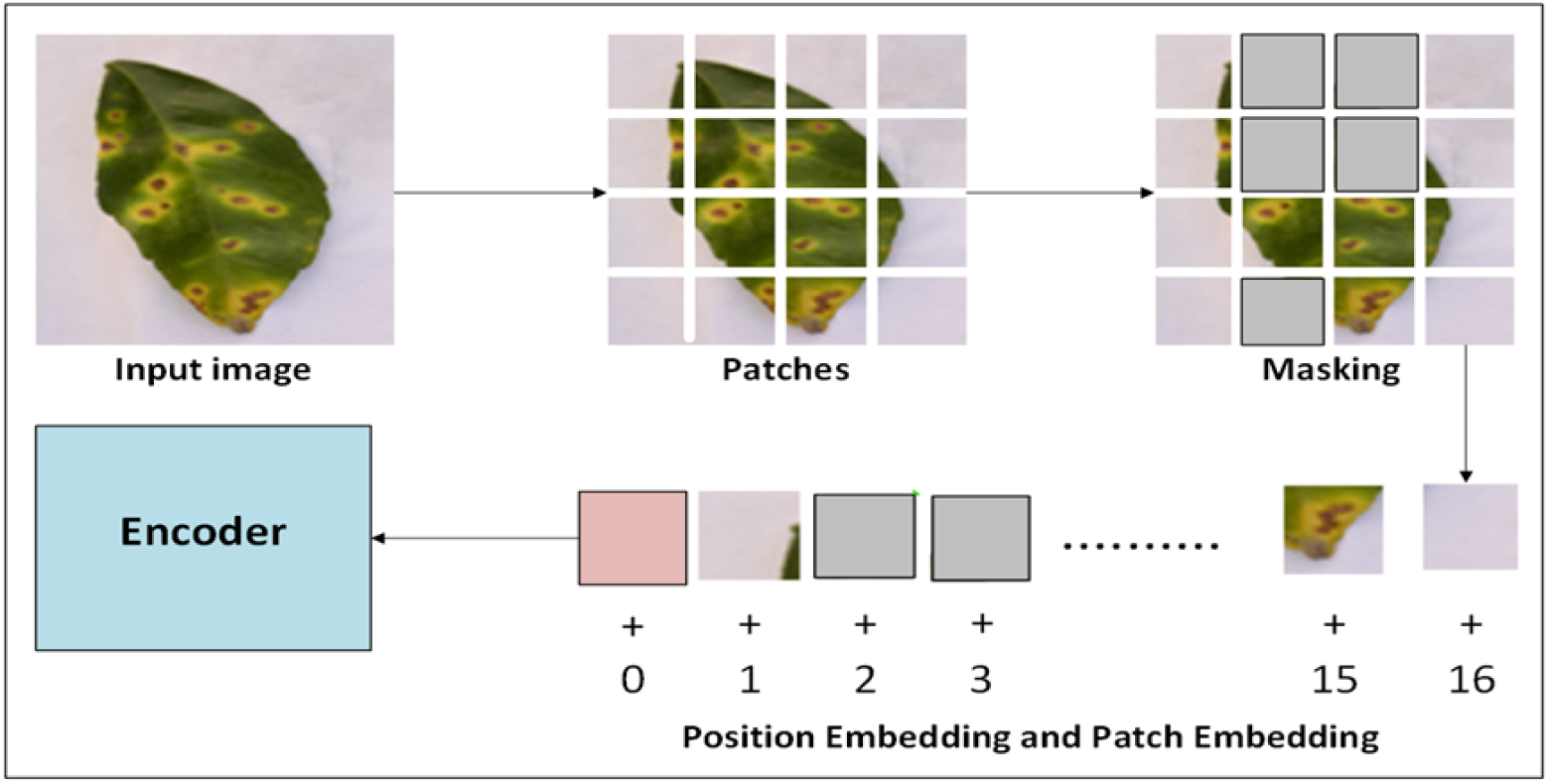
Workflow of BEiT for feature extraction: An input image is split into patches, a random subset of which are masked. The visible patches are then projected into embedding vectors, combined with positional information, and processed by the transformer encoder to generate a contextualized feature representation.

This feature vector is used for classification tasks. Unlike CNNs that focus on local patterns, BEiT captures long-range dependencies across the image, leading to richer representations. It is pre-trained on large-scale datasets, and BEiT provides generalized visual features that can be fine-tuned or reused for specific tasks with limited data. Its attention mechanism allows analysis of where the model focuses during feature extraction. Using BEiT for feature extraction in citrus leaf disease detection provides a powerful alternative to CNNs by leveraging the transformer architecture’s ability to model global context. The [CLS] token embedding serves as a compact yet informative feature vector, enabling effective classification when combined with appropriate machine learning algorithms. Once features are extracted, they are stored in feature matrices (e.g., NumPy arrays) and labeled with corresponding disease categories for the classification task. This methodology is beneficial for agricultural applications where labeled data is often scarce. The global contextual understanding helps in accurately identifying diseases that manifest through complex patterns across the leaf surface. Furthermore, the extracted feature matrices can be used to train simpler, faster classifiers like Support Vector Machines (SVMs), making the overall system efficient for deployment.

One of the biggest advantages of BEiT is its pretraining method, called Masked Image Modeling (MIM) [39]. This allows the model to learn strong visual representations without needing labeled data. During pretraining, about 40% of the image patches are hidden randomly. The model is trained to predict what those hidden patches should look like based on the visible ones. This forces the model to learn about textures, shapes, and how different parts of an image relate to each other. The pretraining process also uses a visual tokenizer, which converts patches into discrete tokens that are similar to words in language models [40], and a small decoder that reconstructs the original image from these tokens. The visual tokenizer is pretrained on a large dataset using a separate self-supervised learning technique like dVAE. The use of discrete tokens is crucial as it transforms the complex, continuous regression task of predicting pixels into a more manageable classification problem. Once pretraining is complete, the decoder is discarded, and the powerful encoder is fine-tuned on downstream tasks such as image classification or segmentation, demonstrating exceptional performance even with limited labeled data.

### Feature Extraction Using MobileNetV2

This study used MobileNetV2 [41] as an efficient feature extraction backbone for citrus disease classification. The lightweight architecture utilizes inverted residuals and depthwise separable convolutions to maintain accuracy while reducing computational demands, making it ideal for resource-constrained applications. It is designed to be computationally efficient while maintaining high classification performance. Figure 5 illustrates the detailed architecture of MobileNetV2.

**Fig 5.**
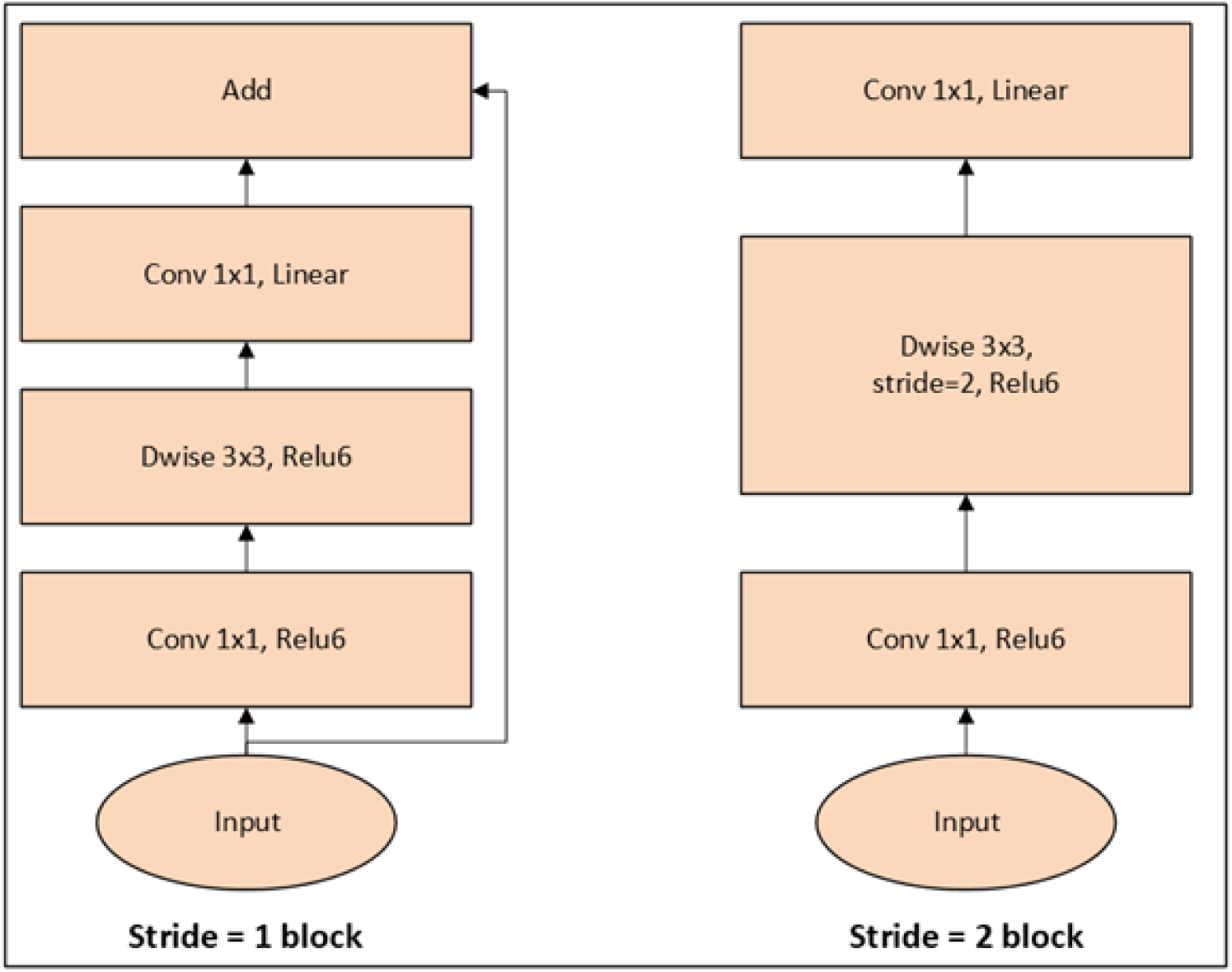
Feature extraction pipeline using MobileNetV2. An input image is processed, and a high-level feature map or vector is extracted from a pre-final layer for downstream tasks.

We utilized a MobileNetV2 model pretrained on the ImageNet dataset to extract generalized visual features from citrus leaf images. The network is truncated before the final classification head, thereby retaining only the feature extraction layers. These layers have been used to capture rich spatial and semantic features, such as leaf texture, shape, and disease-related discolorations. Each input image is first resized to 224×224 pixels and normalized using the ImageNet mean and standard deviation values to match the conditions under which the original MobileNetV2 was trained. This preprocessing step ensures compatibility with the pretrained weights and helps the model generalize well to citrus leaf images. The normalized image is then passed through the convolutional layers of MobileNetV2, and the resulting feature map is flattened to a one-dimensional vector using global average pooling, yielding a fixed-length feature vector of size 1280.

This 1280-dimensional feature vector is used as input to conventional machine learning classifiers such as Support Vector Machines (SVM), Random Forests (RF), or K-Nearest Neighbors (KNN). These classifiers are trained to discriminate between healthy and diseased leaf samples based on the abstract visual features extracted by MobileNetV2. Furthermore, using MobileNetV2 as a feature extractor avoids the need for fine-tuning the entire deep network, reducing both computational cost and overfitting risk. This hybrid approach also facilitates experimentation with different classifiers without retraining the feature extraction backbone.

## Results

The experimental results demonstrate the effectiveness of our proposed hybrid feature extraction approach, which combines BEiT’s global contextual representations with MobileNetV2’s localized features for citrus disease classification. A comprehensive evaluation of four classifiers trained on the combined feature vectors is presented in Table 2.

**Table 1.** Comparative Analysis of existing methodologies for Citrus Disease Detection.

| Ref No. | Method/Architecture | Food/Crop/Fruit | Accuracy |
| --- | --- | --- | --- |
| [7] | Two-stage deep CNN | Citrus leaves | 94.37% |
| [8] | 4-block CNN architecture | Citrus (Blackspot, Canker, Greening) | 96.00% |
| [10] | AlexNet, VGG19 | Citrus leaves | 94.00% |
| [11] | AlexNet, ResNet | Citrus leaves | 97.92% |
| [12] | DenseNet 121 | Citrus leaves | 92.00% |
| [13] | Feature fusion + Whale Optimization Algorithm (WOA) | Citrus leaves | 95.7% |
| [14] | MobileNet, Self-structured classifier | Citrus leaves | 98.00% |
| [15] | Image processing + deep learning | Citrus leaves | 90.00% |
| [16] | CNN (Feature Extractor) + Random Forest | Citrus leaves | 87.00% |
| [17] | Patch-based classification network | Citrus leaves | 95.04% |
| [18] | BridgeNet-19 | Citrus leaves | 95.47% |
| [19] | CNN + Learning Vector Quantization (LVQ) | Citrus leaves | 96.33% |
| [20] | YOLO BP + CSPDarknet53, PANet, Bi-PANet | Citrus (Green) | 86.00% |
| [21] | MobileNetv2 + SVM kernels | Vine leaves | 97.60% |
| [22] | Pretrained CNN + Random Forest | Citrus leaves | 91.66% |
| [24] | Ensemble CNN | Citrus leaves | 96.00% |
| [25] | CNN + LSTM | Citrus (Fruits & Leaves) | 96.00% |
| [26] | CNN-K (CNN + K-means clustering) | Citrus fruit | 99.20% |

**Table 2.**
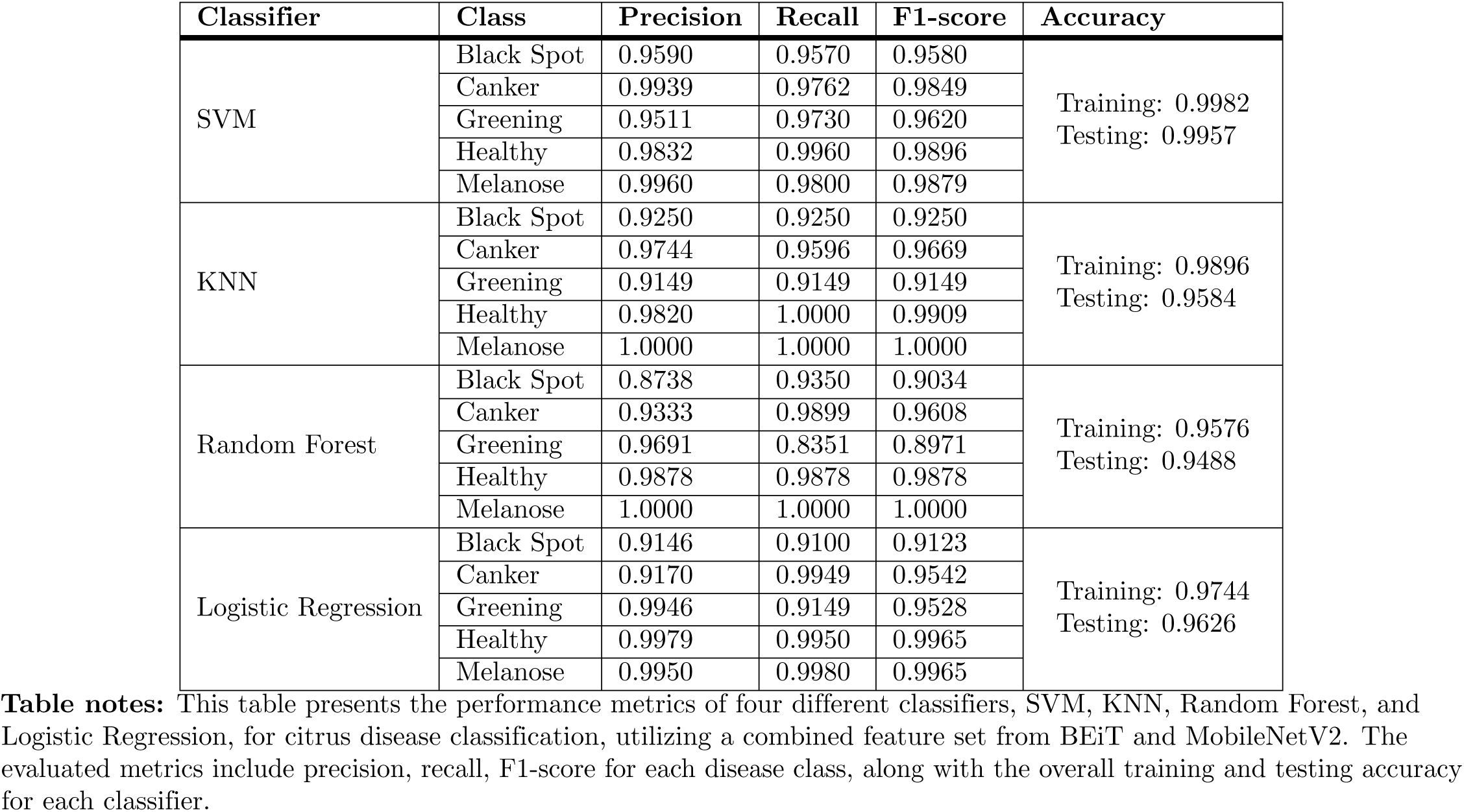
Performance Comparison of Machine Learning Classifiers for Citrus Disease Detection Using a Combined BEiT and MobileNetV2 Feature Set.

| Classifier | Class | Precision | Recall | F1-score | Accuracy |
| --- | --- | --- | --- | --- | --- |
| SVM | Black Spot | 0.9590 | 0.9570 | 0.9580 | Training: 0.9982<br>Testing: 0.9957 |
|  | Canker | 0.9939 | 0.9762 | 0.9849 |  |
|  | Greening | 0.9511 | 0.9730 | 0.9620 |  |
|  | Healthy | 0.9832 | 0.9960 | 0.9896 |  |
|  | Melanose | 0.9960 | 0.9800 | 0.9879 |  |
| KNN | Black Spot | 0.9250 | 0.9250 | 0.9250 | Training: 0.9896<br>Testing: 0.9584 |
|  | Canker | 0.9744 | 0.9596 | 0.9669 |  |
|  | Greening | 0.9149 | 0.9149 | 0.9149 |  |
|  | Healthy | 0.9820 | 1.0000 | 0.9909 |  |
|  | Melanose | 1.0000 | 1.0000 | 1.0000 |  |
| Random Forest | Black Spot | 0.8738 | 0.9350 | 0.9034 | Training: 0.9576<br>Testing: 0.9488 |
|  | Canker | 0.9333 | 0.9899 | 0.9608 |  |
|  | Greening | 0.9691 | 0.8351 | 0.8971 |  |
|  | Healthy | 0.9878 | 0.9878 | 0.9878 |  |
|  | Melanose | 1.0000 | 1.0000 | 1.0000 |  |
| Logistic Regression | Black Spot | 0.9146 | 0.9100 | 0.9123 | Training: 0.9744<br>Testing: 0.9626 |
|  | Canker | 0.9170 | 0.9949 | 0.9542 |  |
|  | Greening | 0.9946 | 0.9149 | 0.9528 |  |
|  | Healthy | 0.9979 | 0.9950 | 0.9965 |  |
|  | Melanose | 0.9950 | 0.9980 | 0.9965 |  |

**Table notes:** This table presents the performance metrics of four different classifiers, SVM, KNN, Random Forest, and Logistic Regression, for citrus disease classification, utilizing a combined feature set from BEiT and MobileNetV2. The evaluated metrics include precision, recall, F1-score for each disease class, along with the overall training and testing accuracy for each classifier.

Among the evaluated models, the Support Vector Machine (SVM) classifier delivered the strongest overall performance, achieving the highest accuracy scores of 99.82% on the training set and 99.57% on the testing set. It maintained high precision, recall, and F1-scores across all disease categories. Performance was particularly strong for Melanose (Precision: 0.9960, Recall: 0.9800, F1-score: 0.9879) and Healthy leaves (Precision: 0.9832, Recall: 0.9960, F1-score: 0.9896). For Black Spot (F1-score: 0.9580) and Greening (F1-score: 0.9620), performance remained robust, though slightly lower, likely due to visual similarities between these diseases. The Canker class also performed well, with an F1-score of 0.9849.

The K-Nearest Neighbors (KNN) classifier achieved a testing accuracy of 95.84% (training accuracy: 98.96%), performing perfectly on Melanose (Precision: 1.0000, Recall: 1.0000, F1-score: 1.0000) and very well on Healthy leaves (F1-score: 0.9909). However, it showed a decline in distinguishing Black Spot (F1-score: 0.9250) and Greening (F1-score: 0.9149).

The Random Forest classifier attained a testing accuracy of 94.88% (training accuracy: 95.76%), with perfect classification for Melanose (F1-score: 1.0000) and near-perfect results for Healthy leaves (F1-score: 0.9878). While it performed well on Canker (F1-score: 0.9608), its ability to classify Black Spot (F1-score: 0.9034) and Greening (F1-score: 0.8971) was weaker.

Logistic Regression achieved a testing accuracy of 96.26% (training accuracy: 97.44%), excelling in classifying Healthy (F1-score: 0.9965) and Melanose (F1-score: 0.9965) leaves. However, it faced difficulties with Black Spot (F1-score: 0.9123), while its performance on Greening was stronger (F1-score: 0.9528).

All experiments were conducted on Google Colab with GPU support, ensuring efficient training and evaluation. The results confirm that SVM is the most effective classifier for this task, delivering consistently high accuracy and balanced performance across all disease categories. This makes our hybrid BEiT-MobileNetV2 approach a practical and reliable solution for automated citrus disease detection in real-world agricultural applications.

The confusion matrices of the top two classifiers, SVM (linear kernel) and Logistic Regression, shown in Figure 6, provide further insight into their performance. The SVM model demonstrates strong classification accuracy, with most classes correctly predicted at very high rates. It shows minimal misclassification, particularly excelling in distinguishing Canker (99.8%), Healthy (99.9%), and Melanose (99.8%). A slight overlap occurs between Greening and Black Spot, with Greening being misclassified in 3.0% of cases, but the model maintains robust overall performance. In contrast, Logistic Regression struggles with certain classes, particularly Black Spot, which is frequently misclassified. While it performs well on Canker (99.5%), Healthy (99.5%), and Melanose (99.8%), its accuracy drops noticeably for Greening, where confusion with Black Spot is evident. These results reinforce that SVM is the more reliable classifier for this task, handling inter-class variations more effectively than Logistic Regression, especially in distinguishing between visually or symptomatically similar diseases such as Black Spot and Greening.

**Fig 6.**
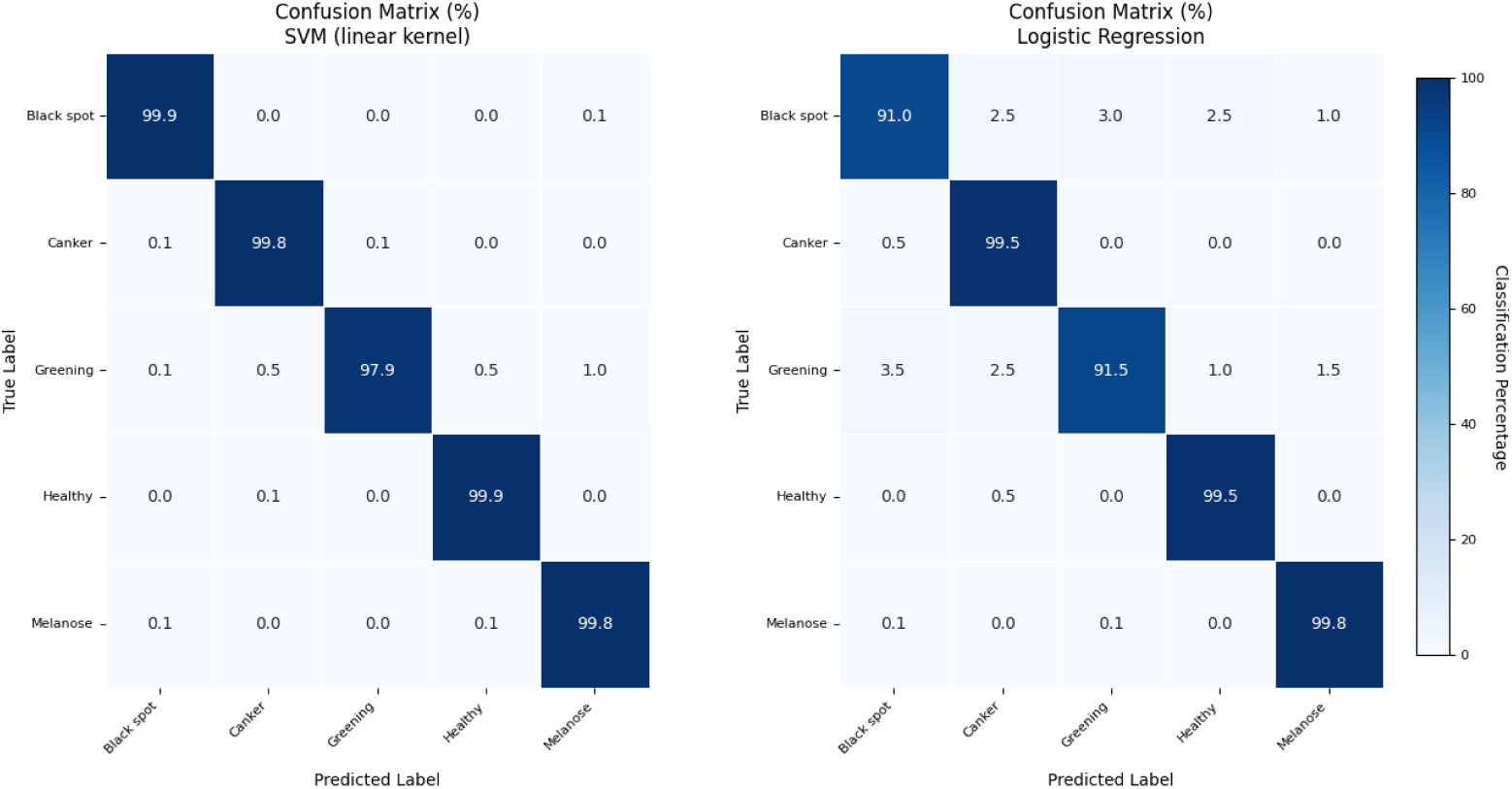
Confusion matrices for the top-performing classifiers: (a) Support Vector Machine (Linear Kernel) and (b) Logistic Regression, illustrating their performance on the citrus disease dataset using the hybrid BEiT and MobileNetV2 features.

### Ablation Study

The ablation study evaluates two feature extraction strategies; transformer-based BEiT and CNN-based MobileNetV2 across four classifiers for citrus leaf disease classification. Overall, the results confirm that BEiT features combined with an SVM classifier deliver the strongest performance, achieving the highest testing accuracy of 98.51%, substantially outperforming the remaining classifier configurations as displayed in Table 3. This combination is especially effective for Canker (F1-score: 0.9849) and Healthy leaves (F1-score: 0.9896), reflecting BEiT’s strong ability to capture global disease structures through its self-attention mechanisms. Performance on Greening is also robust (F1-score: 0.9630), demonstrating BEiT’s reliability even for challenging patterns.

**Table 3.** Performance Comparison of Machine Learning Classifiers for Citrus Disease Detection with BEiT Features.

| Classifier | Class | Precision | Recall | F1-score | Accuracy |
| --- | --- | --- | --- | --- | --- |
| SVM | Black Spot | 0.9590 | 0.9570 | 0.9580 | Training: 0.9921<br>Testing: 0.9851 |
|  | Canker | 0.9939 | 0.9760 | 0.9849 |  |
|  | Greening | 0.9511 | 0.9730 | 0.9630 |  |
|  | Healthy | 0.9832 | 0.9960 | 0.9896 |  |
|  | Melanose | 0.9960 | 0.9800 | 0.9879 |  |
| KNN | Black Spot | 0.9242 | 0.9150 | 0.9196 | Training: 0.9853<br>Testing: 0.9573 |
|  | Canker | 0.9694 | 0.9596 | 0.9645 |  |
|  | Greening | 0.9153 | 0.9202 | 0.9178 |  |
|  | Healthy | 0.9820 | 1.0000 | 0.9909 |  |
|  | Melanose | 1.0000 | 0.9820 | 0.9909 |  |
| Random Forest | Black Spot | 0.8599 | 0.8900 | 0.8747 | Training: 0.9417<br>Testing: 0.9285 |
|  | Canker | 0.9139 | 0.9646 | 0.9386 |  |
|  | Greening | 0.9157 | 0.8085 | 0.8588 |  |
|  | Healthy | 0.9643 | 0.9878 | 0.9759 |  |
|  | Melanose | 0.9712 | 0.9804 | 0.9707 |  |
| Logistic Regression | Black Spot | 0.8657 | 0.8650 | 0.8650 | Training: 0.9594<br>Testing: 0.9413 |
|  | Canker | 0.9697 | 0.9900 | 0.9798 |  |
|  | Greening | 0.8738 | 0.8705 | 0.8721 |  |
|  | Healthy | 0.9849 | 0.9800 | 0.9824 |  |
|  | Melanose | 0.9939 | 0.9830 | 0.9884 |  |

**Table notes:** This table presents the performance metrics of four different classifiers (SVM, KNN, Random Forest, and Logistic Regression) for citrus disease classification using features extracted from the BEiT model. The metrics include precision, recall, F1-score for each disease class, and the overall training and testing accuracy for each classifier.

Among the alternative classifiers using BEiT features, KNN achieves a respectable testing accuracy of 95.73%, but shows reduced performance in visually similar disease types such as Black Spot (F1-score: 0.9196). Logistic Regression, with a testing accuracy of 94.13%, performs well on Canker (F1-score: 0.9798) and Melanose (F1-score: 0.9884), yet continues to struggle with Greening (F1-score: 0.8721). Random Forest yields the lowest testing accuracy (92.85%), with its weakest result observed for Greening (F1-score: 0.8588), likely due to its dependence on localized decision boundaries that fail to capture broad spatial disease cues.

These findings reinforce that BEiT’s ability to model long-range dependencies produces richer and more discriminative feature embeddings than conventional CNN-based methods, particularly for diseases characterized by irregular or diffused patterns. At the same time, the choice of classifier remains crucial. SVM’s strength in constructing optimal decision boundaries in high-dimensional feature spaces makes it especially compatible with BEiT embeddings, allowing it to fully exploit global feature representations. Simpler classifiers like KNN and Logistic Regression offer computational advantages but exhibit larger drops in performance for classes with subtle lesion characteristics, indicating that robustness requires both advanced feature extraction and an equally capable classification algorithm.

Finally, although BEiT demonstrates excellent classification performance, its effectiveness decreases when not supported by deep learning–derived features. For instance, the precision for Black Spot (0.9590)—while high—shows that purely global representations may still overlook fine-grained local texture variations that CNNs naturally detect. This highlights the complementary nature of global transformer features and local CNN descriptors, emphasizing the potential benefit of hybrid or fused approaches for comprehensive citrus disease recognition.

Figure 7 presents the confusion matrices for the SVM and Logistic Regression classifiers using only BEiT features. The SVM classifier demonstrates strong overall performance, achieving very high class-wise accuracies, with exceptionally accurate predictions for Healthy (99.4%), Canker (97.6%), and Greening (97.3%). However, small but important misclassifications remain, particularly the confusion between Black Spot and Greening, and occasional mixing between Canker and Healthy. Logistic Regression shows a more noticeable decline in accuracy, especially for Black Spot, where accuracy drops to 86.5%, with frequent misclassification into Greening, Healthy, and Canker. Greening is similarly affected, with Logistic Regression achieving only 86.8%, reflecting substantial confusion with Black Spot and Melanose. Despite this, both models maintain strong classification on Melanose, with accuracies above 98%. Overall, the increased inter-class confusion—more pronounced in Logistic Regression—indicates that BEiT’s global features alone do not fully capture the fine-grained visual differences between diseases such as Black Spot, Greening, and Canker. This further supports the benefit of integrating complementary local feature extractors like MobileNetV2 to enhance discriminability.

**Fig 7.**
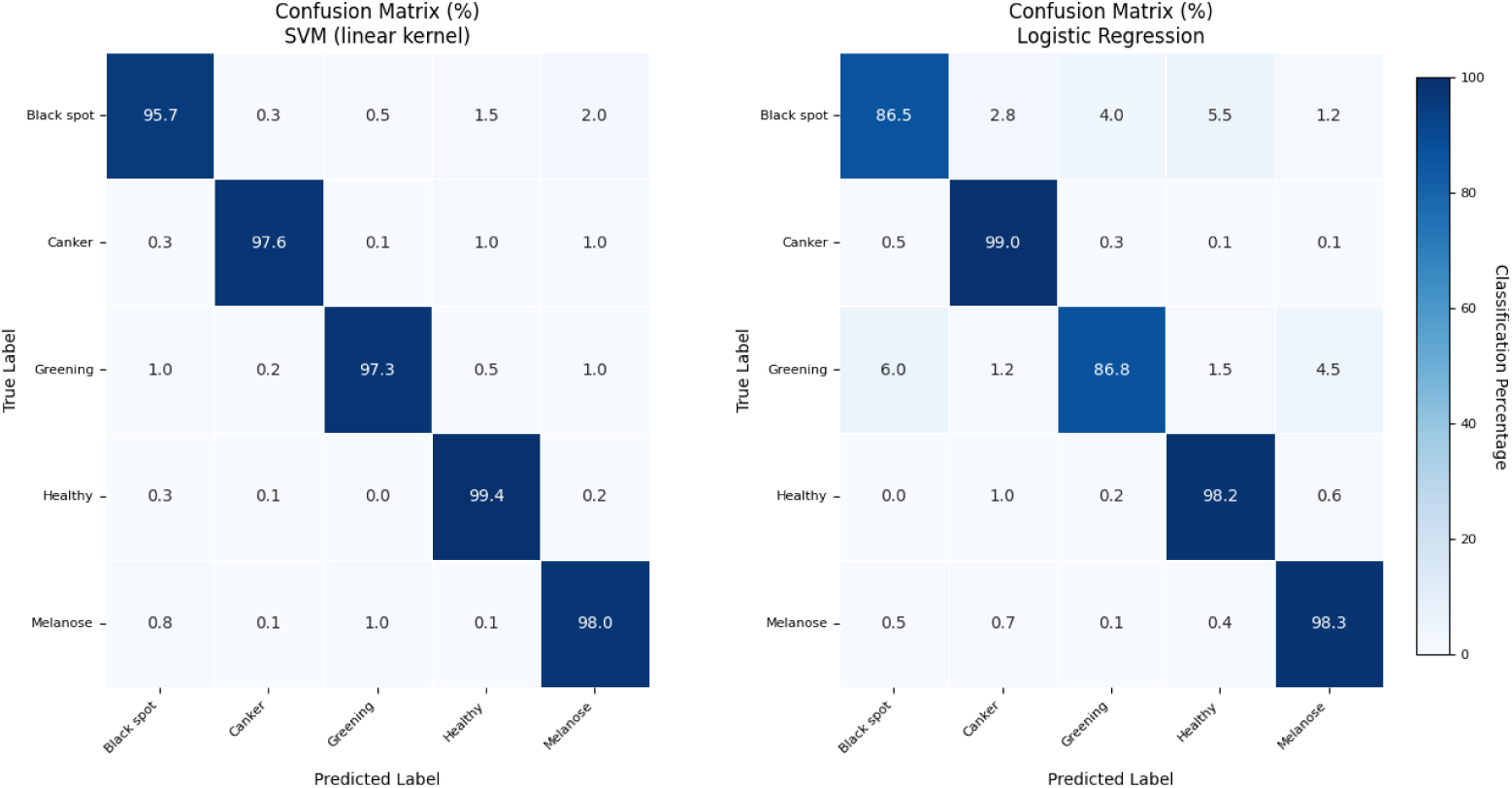
Confusion matrices for the (a) SVM and (b) Logistic Regression classifiers trained exclusively on BEiT-derived features, revealing specific inter-class confusion patterns, particularly between Greening and Black Spot.

The CNN-based MobileNetV2 features exhibit comparatively reduced performance when used independently, as reflected in the results summarized in Table 4. Among the four classifiers, SVM remains the strongest performer, achieving a testing accuracy of 95.09%, though this marks a noticeable drop from the 98.51% achieved using BEiT features alone. SVM delivers high effectiveness on Canker (F1: 0.9783) and Melanose (F1: 0.9772), yet its performance declines for Black Spot (F1: 0.8723), confirming that CNN features struggle with diseases requiring broader contextual understanding.

**Table 4.** Performance Comparison of Different Classifiers for Citrus Disease Detection Using MobileNetV2 Features.

| Classifier | Class | Precision | Recall | F1-score | Accuracy |
| --- | --- | --- | --- | --- | --- |
| SVM | Black Spot | 0.8342 | 0.9130 | 0.8723 | Training: 0.9560<br>Testing: 0.9509 |
|  | Canker | 0.9783 | 0.9781 | 0.9783 |  |
|  | Greening | 0.9464 | 0.9457 | 0.9456 |  |
|  | Healthy | 0.9500 | 0.9500 | 0.9500 |  |
|  | Melanose | 0.9772 | 0.9773 | 0.9772 |  |
| KNN | Black Spot | 0.9171 | 0.9400 | 0.9284 | Training: 0.9887<br>Testing: 0.9605 |
|  | Canker | 0.9694 | 0.9596 | 0.9645 |  |
|  | Greening | 0.9344 | 0.9096 | 0.9218 |  |
|  | Healthy | 0.9880 | 1.0000 | 0.9939 |  |
|  | Melanose | 1.0000 | 1.0000 | 1.0000 |  |
| Random Forest | Black Spot | 0.7931 | 0.8050 | 0.7990 | Training: 0.8819<br>Testing: 0.8634 |
|  | Canker | 0.7974 | 0.9343 | 0.8605 |  |
|  | Greening | 0.8819 | 0.6755 | 0.7651 |  |
|  | Healthy | 0.8713 | 0.9085 | 0.8896 |  |
|  | Melanose | 1.0000 | 1.0000 | 1.0000 |  |
| Logistic Regression | Black Spot | 0.8500 | 0.7880 | 0.8178 | Training: 0.9173<br>Testing: 0.9168 |
|  | Canker | 0.9800 | 0.9790 | 0.9795 |  |
|  | Greening | 0.9733 | 0.8365 | 0.8990 |  |
|  | Healthy | 0.9910 | 0.9939 | 0.9924 |  |
|  | Melanose | 1.0000 | 0.9780 | 0.9889 |  |

**Table notes:** This table presents the performance metrics of four different classifiers (SVM, KNN, Random Forest, and Logistic Regression) for citrus disease classification using features extracted from MobileNetV2. The evaluated metrics include precision, recall, and F1-score for each disease class, along with the overall training and testing accuracy for each classifier.

KNN performs slightly better than SVM for certain classes, reaching the highest MobileNetV2 testing accuracy of 96.05%, supported by perfect detection of Melanose (F1: 1.0000) and strong identification of Healthy leaves (F1: 0.9939). However, its performance weakens for Greening (F1: 0.9218), indicating limited capability in distinguishing diseases with subtle inter-class variations.

Random Forest shows the greatest decline in performance, achieving a testing accuracy of only 86.34%, with notable confusion for Greening (F1: 0.7651) and Black Spot (F1: 0.7990). Logistic Regression also underperforms relative to SVM and KNN, obtaining a testing accuracy of 91.68%, and showing reduced effectiveness for Black Spot (F1: 0.8178) and Greening (F1: 0.8990) despite strong results for Canker and Melanose.

Collectively, these results show that MobileNetV2 alone yields 3–5% lower accuracy compared with BEiT across classifiers, emphasizing the limitations of CNNs in capturing global disease patterns. While MobileNetV2 excels at recognizing localized textures—evident in the consistently high Melanose scores—its standalone performance weakens for diseases requiring global structural understanding, such as Black Spot and Greening. This reinforces that hybrid feature extraction is essential: BEiT’s global attention mechanisms compensate for CNN’s locality bias, while MobileNetV2 contributes fine-grained spatial details that enrich BEiT’s features. Together, they offer a more comprehensive disease representation than either architecture can achieve alone.

The confusion matrices in Figure 8 present the classification performance when using only MobileNetV2 features. The SVM classifier demonstrates reasonably strong behavior overall, particularly for Canker (97.8%) and Melanose (97.7%), with very few misclassifications in these categories. Black Spot is also recognized well (91.5% accuracy), although small confusions with Canker (2.7%) and Greening (1.8%) are present. Greening shows comparatively lower accuracy (84.6%), with notable confusion toward Black Spot (3.2%) and Healthy (5.8%). Healthy leaves are identified with 95% accuracy, with minor misclassification into Melanose (2.5%). In contrast, Logistic Regression performs less effectively across several classes. While it still classifies Canker (98.0%) and Melanose (97.8%) very well, its performance for Black Spot drops to 85%, with increased confusion toward Greening (4.2%) and Healthy (5.8%). Greening is the most challenging class for Logistic Regression, achieving only 76.6% accuracy, with a noticeable proportion misclassified as Melanose (7.9%) and Black Spot (6.8%). Healthy leaves, however, are recognized reliably with 98.5% accuracy, showing only minimal confusion across other classes. Overall, these results indicate that MobileNetV2 features alone capture local texture information well—evident from the strong performance on Canker and Melanose—but they struggle with diseases that share overlapping visual cues, such as Greening and Black Spot. Logistic Regression, in particular, shows greater sensitivity to these feature limitations. This reinforces the need for complementary transformer-based global representations (e.g., BEiT) to improve separability among visually similar classes and achieve more balanced disease characterization.

**Fig 8.**
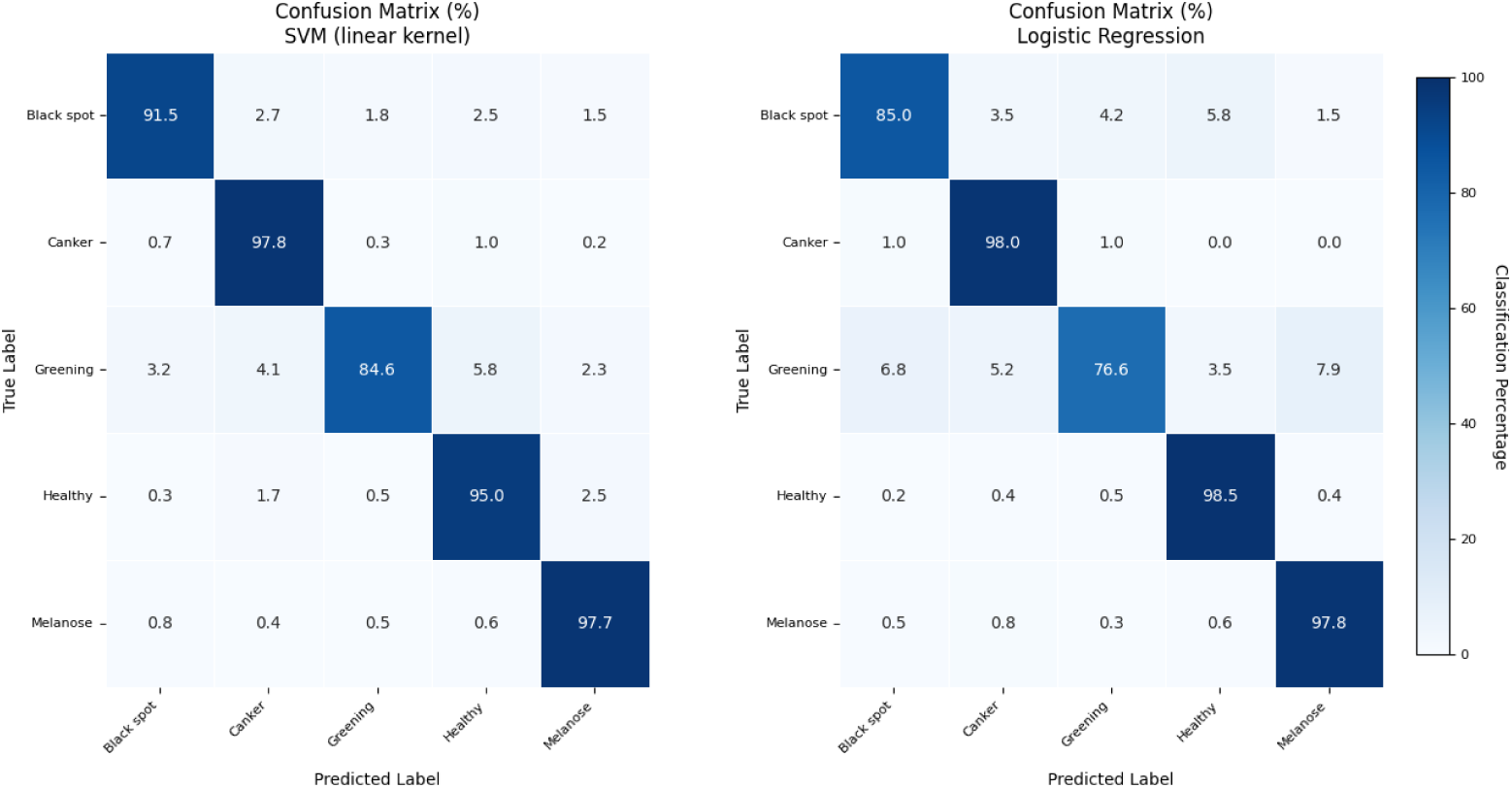
Performance of classifiers with MobileNetV2 features, highlighting the architectural limitation of CNNs in capturing global context.

Our results and comparative analysis demonstrate that while both feature extraction methods achieve competent performance for citrus disease classification, BEiT’s transformer-based approach consistently outperforms MobileNetV2’s CNN architecture, particularly in distinguishing visually similar diseases. The superiority of BEiT stems from its ability to capture nuanced pathological patterns through global contextual analysis, whereas MobileNetV2 provides a more resource-efficient alternative with slightly reduced accuracy. SVM emerges as the most effective classifier across both feature types, reliably leveraging their respective strengths. These findings suggest that the choice between methods should consider application requirements: BEiT for maximum diagnostic precision in controlled environments, and MobileNetV2 for scalable deployments where computational efficiency is prioritized. The results collectively highlight the importance of feature representation in plant disease analysis while validating the viability of both approaches for agricultural applications.

Even though the BEiT-SVM combination shows higher performance than the other models, we do recognize two important challenges to deploying the model: First, the computational requirements of the transformer architecture and their inefficiency in resource-constrained agricultural settings. Second, the dataset, while sufficient for proof-of-concept testing, is missing some of the disease symptoms encountered in real-world agricultural scenarios. Future research can explore lightweight transformer architectures that can be deployed in edge devices. Furthermore, future datasets containing a larger variety of disease severities, disease stages and imaging conditions will be essential to improve the generalization of the developed models. Another promising approach is the development of hybrid architectures, combining the highly parameter efficient CNNs with the global modeling capabilities of the transformers, in order to achieve the best balance of efficiency. The aim is to put these advanced diagnostic capabilities into strong, usable diagnostics to provide real value to farmers.

## Conclusion

CitriBEiTNet, our proposed hybrid MobileNetV2-BEiT framework, demonstrates remarkable effectiveness in automating citrus leaf disease diagnosis. By combining MobileNetV2’s efficient local feature extraction with BEiT’s global contextual understanding through self-attention mechanisms, the model achieves superior classification performance compared to standalone deep learning architectures. The framework’s robustness is validated through extensive experiments, where it attained an exceptional testing accuracy. The success of CitriBEiTNet is due to its ability to capture both fine-grained texture patterns and long-range dependencies in diseased leaf images, while maintaining computational efficiency suitable for real-world deployment. The minimal performance gap between training and testing phases confirms the model’s strong generalization capabilities. This research makes significant contributions to precision agriculture by introducing a novel hybrid architecture that effectively bridges convolutional and transformer-based architectures and provides a scalable solution that balances accuracy with computational efficiency. The CitriBEiTNet framework represents a meaningful advancement in AI-driven plant healthcare, with substantial potential to enhance crop productivity and food security. Figure 9 shows the comparison of our model with state-of-the-art methods.

**Fig 9.**
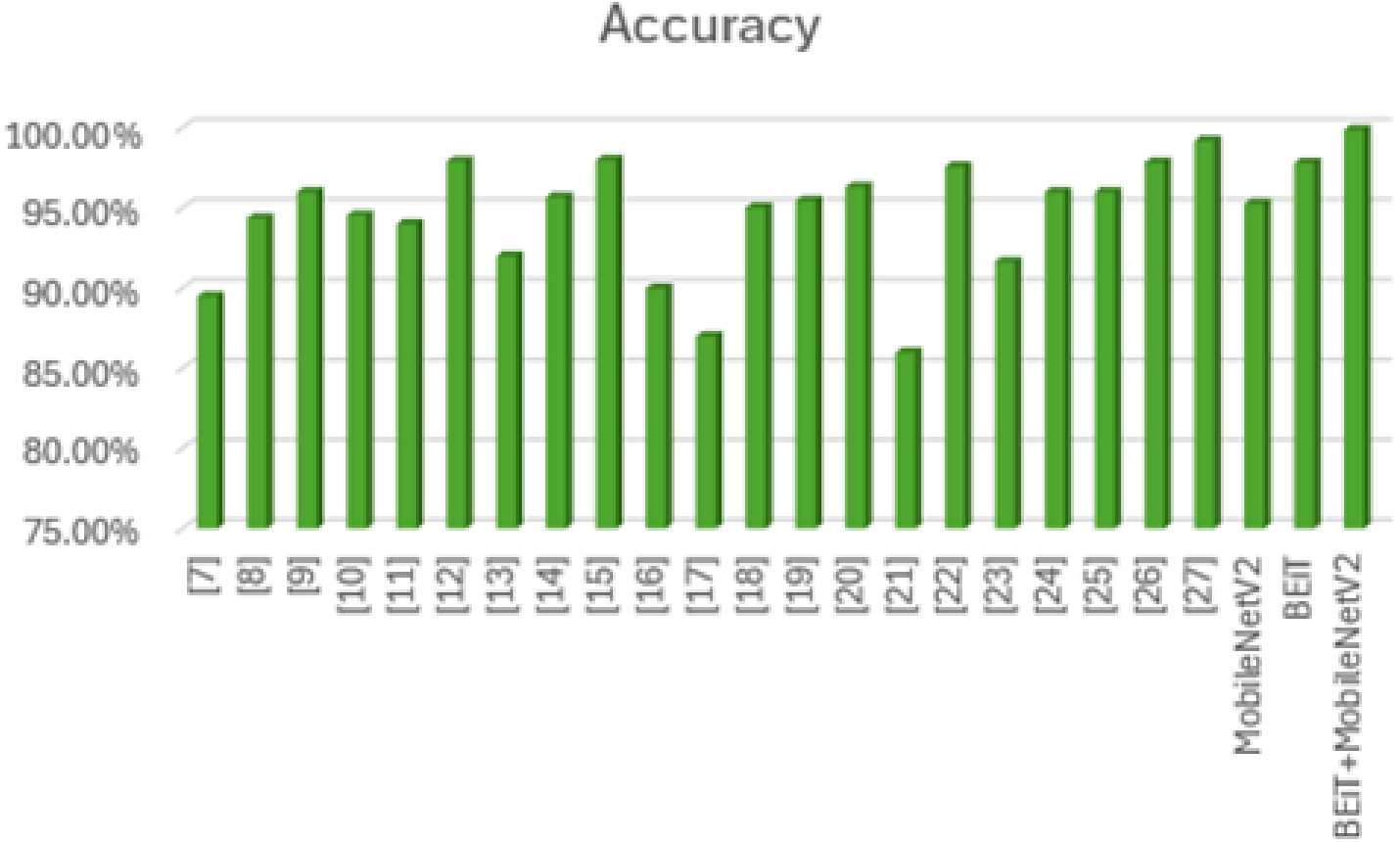
Performance comparison of the proposed hybrid (BEiT-MobileNetV2) model against state-of-the-art methods on the citrus disease classification task.

## Acknowledgments

The Department of Computer Science, University of Engineering and Technology, Taxila facilitated us with the Image Processing Lab and other resources for the experimentation of this research.

